# Vaginal Microbiome Topic Modelling of Laboring Ugandan Women With and Without Fever

**DOI:** 10.1101/2020.12.01.401703

**Authors:** Mercedeh Movassagh, Lisa M. Bebell, Kathy Burgoine, Christine Hehnly, Lijun Zhang, Kim Moran, Kathryn Sheldon, Shamim A. Sinnar, Edith Mbabazi, Elias Kumbakumba, Joel Bazira, Moses Ochoa, Ronnie Mulondo, Brian K. Nsubuga, Andrew Weeks, Melissa Gladstone, Peter Olupot-Olupot, Joseph Ngonzi, Drucilla J. Roberts, Frederick A. Meier, Rafael Irizarry, James Broach, Steven J. Schiff, Joseph N. Paulson

**Affiliations:** Harvard T.H. Chan School of Public Health, Department of Biostatistics, Dana Farber Cancer Institute, Department of Data Sciences; Massachusetts General Hospital, Department of Medicine, Division of Infectious Diseases and Harvard Medical School; Mbale Clinical Research Institute; Institute for Personalized Medicine, Dept. of Biochemistry and Molecular Biology, Penn State University College of Medicine; Institute for Personalized Medicine, Penn State College of Medicine; Pennsylvania State University College of Medicine; CURE Children's Hospital of Uganda; Dept of Microbiology, Mbarara University of Science and Technology; Department of Paediatrics and Child Health, Mbarara University of Science and Technology; Sanyu Research Unit, University of Liverpool, Liverpool Women's Hospital, Crown Street, Liverpool L8 7SS, UK; University of Liverpool; Mbale Clinical Research Institute/Busitema University; Mbarara University of Science and Technology, Faculty of Medicine, Department of Obstetrics and Gynaecology; Massachusetts General Hospital, Department of Pathology; Department of Pathology Wayne State University School of Medicine; Dana Farber Cancer Institue; Penn State College of Medicine; Genentech, Inc.

## Abstract

The composition of the maternal vaginal microbiome may influence the duration of pregnancy, onset of labor and even neonatal outcomes. Maternal microbiome research in sub Saharan-Africa has focused on non-pregnant and postpartum composition of the vaginal microbiome. We examined the vaginal microbiome composition of 99 laboring Ugandan women using routine microbiology and 16S ribosomal DNA sequencing from two hypervariable regions (V1-V2 and V3-V4), using standard hierarchical methods. We then introduce Grades of Membership (GoM) modeling for the vaginal microbiome, a method often used in the text mining machine learning literature. Leveraging GoM models, we create a basis composed of a small number of microbial ‘topic’s whose linear combination optimally represents each patient yielding more accurate associations. We identified relationships between defined communities and the presentation or absence of intrapartum fever. Using a random forest model we showed that by including novel microbial topic models we improved upon clinical variables to predict maternal fever. We also show by integrating clinical variables with a microbial topic model into this model found young maternal age, fever report earlier in the current pregnancy, and longer labors, as well as a more diverse, less *Lactobacillus* dominated microbiome were features of labor associated with intrapartum fever. These results better define relationships between presentation or absence of intrapartum fever, demographics, peripartum course, and vaginal microbial communities, and improve our understanding of the impact of the microbiome on maternal and neonatal infection risk.

## Introduction

The vaginal microbiome consists of an ecological community of microorganisms which are important in both maternal and neonatal health ^1^. For neonates, exposure to the vaginal microbiome during birth or through premature rupture of membranes is an important route to early onset neonatal sepsis ^2^. During pregnancy, the vaginal microbiome composition is known to change, which has a role in ascending infection in puerperal sepsis ^3^. Routine screening and treatment for group B Streptococcus (GBS) has reduced the rate of neonatal GBS infection in high-income countries ^4^. In sub-Saharan Africa (sSA), group B Streptococcal infections are relatively uncommon in early onset neonatal sepsis and there is a lack of understanding of how the peripartum vaginal microbiome contributes to maternal and neonatal disease ^5,6^. Most maternal microbiome research in sSA has focused mainly on non-pregnant or postpartum composition of the vaginal microbiome ^7,8^.

In North America, the composition of the vaginal microbiome in reproductive aged women has been characterized using 16S ribosomal DNA sequencing (predominantly V3-V4 hypervariable regions) to determine the presence of culturable and unculturable bacteria from women of reproductive age ^9^. Vaginal profiles were categorized into five (I-V) distinct bacterial communities by hierarchical clustering where patients were assigned a single community ^9^. Subsequent work revealed the temporal dynamics of the bacterial communities and a large dynamic shift of community composition over a 16-week time period for some patients. This highlighted why a single community assignment was often not stable over time and the need for a more dynamic representation of a patients’ community.

Overall, current studies in African women suggest vaginal community compositions distinct from European and North American women ^10^. Additionally, community associations with HIV status and other diseases such as human papillomavirus (HPV) and vaginal infections have been described ^10,11^. In resource-poor settings in sSA, such as Uganda, there is a higher prevalence of many factors that could potentially affect vaginal microbial diversity, including off-label antibiotic use, sexually transmitted infections (STIs) including HIV and *Chlamydia*, endemic malaria and cytomegalovirus (CMV) infections ^10,12–14^. Furthermore, these comorbidities impact microbial diversity associated with infant pneumonia, acute diarrhea, sepsis and postinfectious hydrocephalus ^14–16^. Though women living in Africa are at greater risk for malaria, sepsis, and infectious diarrhea, the effect of these infections on the vaginal microbiome has not been characterized ^17–20^.

The neonate’s first exposure to microbes is ideally through contact with the maternal vaginal and gut microbiota during the birth process ^21,22^. Maternal vaginal communities change during pregnancy, thought in part to result from hormonal changes ^23^. Higher estrogen levels promote the growth of lactic acid-producing bacteria, shifting the microbiome towards predominantly high-*Lactobacillus* communities or increasing diversity ^23,24^. Previous research has also suggested that children born via cesarean delivery compared to those born vaginally are inherently more susceptible to developing autoimmune diseases due to absence of bacteria such as *Bifidobacterium* in their intestinal microbiome ^25,26^. With GBS neonatal sepsis relatively uncommon in sSA, how the composition of the vaginal microbiome in an intrapartum sSA woman renders her infant more susceptible to neonatal infections remains uncharacterized at this time ^26^.

Most maternal microbiome research in sSA has focused on non-pregnancy or postpartum composition of the vaginal microbiome ^7,8^. Here we sought to comprehensively define the structure of the vaginal microbiome of 99 laboring intrapartum women in Uganda through 16S sequencing of V1-V2 and V3-V4 hypervariable regions. Given the importance of maternal intrapartum fever as an indicator of infection, we incorporated novel modeling methods to more fully characterize the vaginal microbiomes between febrile and afebrile mothers.

## Results

### Clinical characterization

Maternal vaginal samples were obtained from 99 Ugandan intrapartum women during active labor but before birth after obtaining written informed consent. Women were enrolled after presenting to hospital in labor for delivery in two districts in Uganda: Mbarara (n=50) in western Uganda and Mbale (n=49) in eastern Uganda (see <u>Supplementary Materials</u>, <u>Online Methods</u>). Various clinical and microbiological features were collected and assessed comparing maternal fever status (Table 1, <u>Supplemental Table 1</u>, <u>Supplemental Table 2</u> and <u>Supplementary Materials</u>).

**Table 1.** Overview of Clinical Characterization febrile versus afebrile **a**:N=(49,49), **b**:N=(48,49), **c**:N=(46,44), **d**:N=(47,50). IQR stands for interquartile range. Odds ratio for hours in labor was estimated for every 10 hours. Samples site collection was Mbale versus Mbarara.

| Characteristic (N = 99) | *afebrile* | *febrile* | *Odds Ratio (95% CI)* |
| --- | --- | --- | --- |
|  | *Total n = 49* | *Total n = 50* | *Univariate* |
| Average temperature at recruitment (mean, SD) | 36.6 (0.3) | 38.3(0.2) | - |
| Age in years (median, IQR) | 26 (7) | 22 (6.3) | 0.03 (0.00, 0.26) |
| HIV-infected n(%) a | 5 (10%) | 5 (10%) | 1.00 (0.26, 3.83) |
| Parity (mean, SD) | 2.8 (1.9) | 2.3 (2) | 0.61 (0.28,1.32) |
| Self-reported fever in last 7days prior to delivery (%) | 4 (6%) | 25 (50%) | 10.56 (3.56, 39.08) |
| Self-reported vaginal infection in last 1month (%) b | 3 (6%) | 3 (10%) | 1.70 (0.39, 8.71) |
| Intrapartum antimicrobial use (%) c | 2 (4%) | 5 (11%) | 2.82 (0.57, 20.47) |
| Hours in labor (median, IQR) | 18 (38) | 33 (30) | 1.19(0.98,1.46) |
| Cesarean delivery (%) | 11 (22%) | 24 (48%) | 3.19 (1.36, 7.84) |
| Malaria RDT or blood smear positive in labor(%) d | 4 (8%) | 16 (32%) | 5.05 (1.68, 18.92) |
| Sample collection site (%) | 24 (48%) | 25 (50%) | 1.04 (0.47, 2.30) |
| Maternal CMV viral load (%) | 16 (32%) | 17 (34%) | 1.06 (0.45, 2.47) |

### Vaginal microbiome communities of afebrile and febrile laboring Ugandan women

An overview of our 16S ribosomal sequencing approach on the V1-V2 and V3-V4 hypervariable regions and taxonomic assignment method is provided in Fig. 1a and in the <u>Supplementary Materials</u>. We assessed bacterial recovery in culture and compared it to 16S results (see <u>Supplementary Materials</u>).

**Fig. 1:**
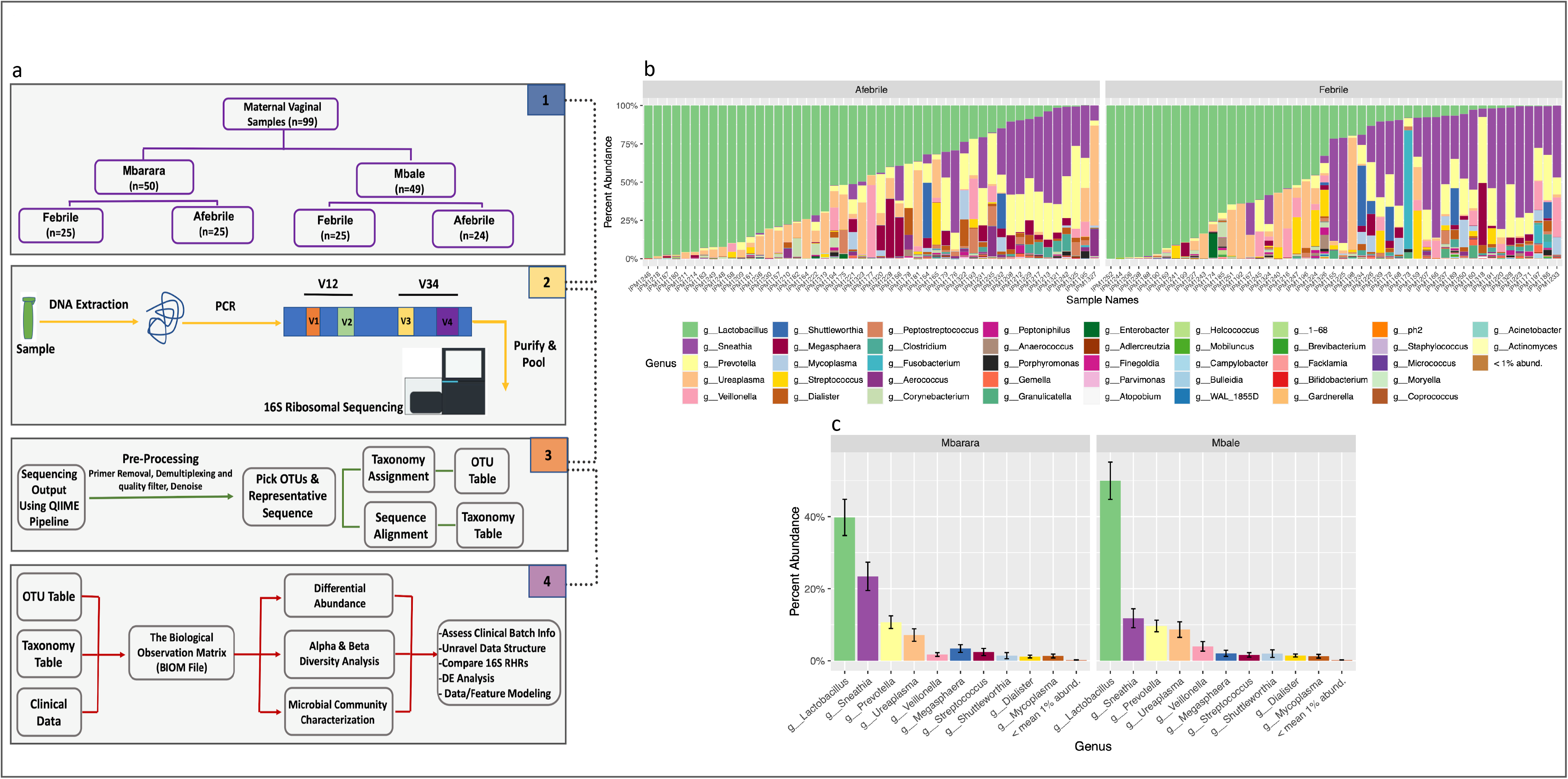
Overall pipeline and structure for 16S ribosomal sequencing of maternal vaginal samples. **a**, Maternal vaginal samples were collected from two hospital sites in Uganda (Mbarara and Mbale) and categorized by intrapartum fever status. DNA was extracted and samples underwent library preparation and sequencing on two ribosomal hypervariable regions V1-V2 and V3-V4. The sequence output was pre-processed utilizing the QIIME1 pipeline (Methods) and samples were further processed for downstream differential abundance (DA) and modeling in concordance with various clinical and technical variables. **b,** Percentage abundance of bacteria on the genus level based on the febrile status of samples. **c,** Mean percent abundance of bacteria (agglomerated on the genus level) by enrollment site. (g_) denotes the bacteria naming based on genus taxonomic level.

We aimed to dissect patterns within our data which could unravel the relationship between vaginal microbiome structure and maternal fever. Through taxonomic assignment and downstream analysis, we estimated the percent abundance of top genera represented in our dataset Fig. 1b. As expected, *Lactobacillus* was the most predominant genus in the vaginal microbiome of laboring women ^23^. *Sneathia*, often associated with BV, was the second most predominant genus ^10^, followed by *Ureaplasma* and *Prevotella,* all of which are normal commensal vaginal flora ^27,28^. The distribution of the top genera of bacteria identified was primarily consistent across both sample collection sites Fig. 1c.

We initially examined the vaginal communities through previously described hierarchical clustering methods. Since there has not been a consensus on the number of communities in the vaginal microbiome of African women, we assumed agreement with the number of communities expected in the European and African-American samples and picked five as the initial number of clusters to investigate ^10^. We pursued two approaches; the traditional hierarchical approach (see <u>Supplementary Materials</u>) and a modification that incorporates differential abundance analysis.

We refined sample clusters on a subset of the taxa differentially associated with the traditional hierarchical clustering to identify communities on taxa that could be used as a biomarker in resolving community membership and filtering taxa that are not differentially abundant between communities, <u>Online Methods</u>. Community (CMT) 1 was identified as the *Lactobacillus* rich (minus *L. iners*) community (<u>Supplemental Table 3</u>). CMT 2 was mainly composed of high *Lactobacillus iners* species, while CMT 3 appeared to be a similar community, while containing a greater abundance of *Finegoldia*, *Veillonella* and *Streptococcus* genera. CMTs 4a and 4b displayed more diverse communities with high levels of *Sneathia, Diallister, Gemella, Clostridium* and *Prevotella* genera. CMT 4b had higher levels of *Parvimonas, Clostridium, Mycoplasma, Adlercreutzia* and *Mycoplasma* genera in comparison to CMT 4a. We additionally identified unique bacterial genera including *Adlercreutzia, Granulicatella, Bulledia, Staphylococcus, Micrococcus,* and *Fusobacterium* in this cohort and community (<u>Supplemental Table 3</u>). An association between the maternal CMV viral load and the vaginal microbial community was not observed (P = 0.57). Furthermore, there was no association between maternal intrapartum fever and community classification (P = 0.52) Fig. 2. When classified using the V1-V2 hypervariable region, similar communities were formed; however, this region was not as efficient at speciating *Lactobacillus* species, but identified other unique species compared to the V3-V4 hypervariable region ^29^ <u>Supplemental Figure 2</u>, <u>Supplemental Table 2</u>.

**Fig. 2:**
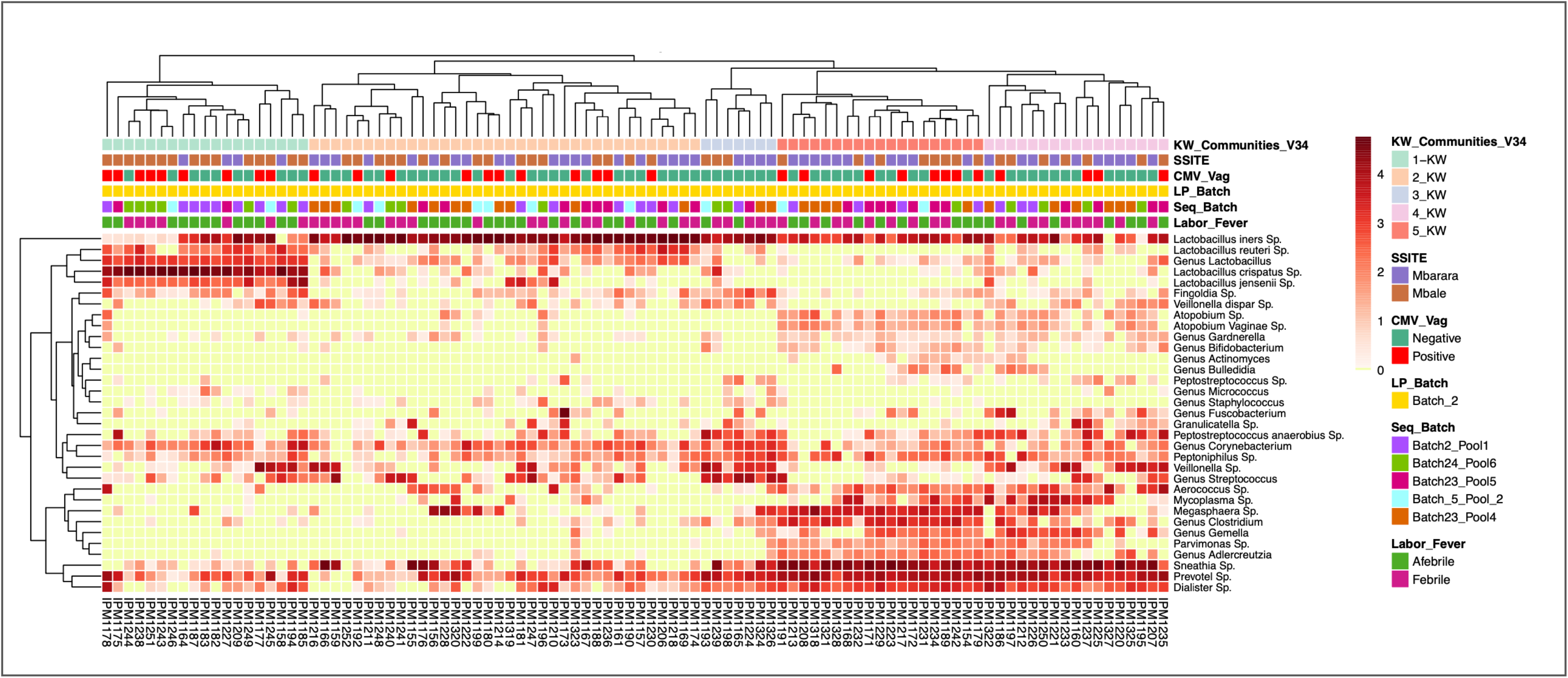
Vaginal bacterial community characterization heatmaps of intrapartum Ugandan women through hierarchical clustering. Vaginal bacterial community classification through selected bacteria after Kruskal-Wallis (KW) test. KW_Communities_V34, are the communities identified by bacteria selected from KW test. Color of the heatmap represents log10 normalized counts of species, yellow represents zero counts. Annotations are as follows: KW_Communities_V34 is the vaginal community identified from V3-V4 regions through hierarchical clustering followed by bacteria selected by KW test. CMV_Vag, represents CMV status of the vaginal samples identified by PCR, LP_Batch is the library preparation batch, Seq_Batch is the sequencing batch which the samples were processed in, Labor_Fever is the fever status of the laboring woman (see Methods for definitions, SSITE is the sample collection site (Mbarara or Mbale).

We assessed alpha diversity (Shannon and Simpson) ^30^ of the vaginal microbiome comparing fever and CMT states across our samples. We observed no significant difference in alpha or beta diversity using either V1-V2 or V3-V4 between febrile states (P =0.25, P =0.12) Fig. 3a and 3c. On the contrary, the microbial communities assigned from hierarchical clustering significantly explained the alpha diversity (Shannon P < 1.78e-08, Simpson P < 3.91e-04) Fig. 3b and 3d and Supplemental Fig. 3a, b, c, d. Overall we find that the structure of our sample cohort assessed by alpha and beta diversity was highly driven by microbial communities assigned by hierarchical clustering and not by maternal fever status.

**Fig. 3:**
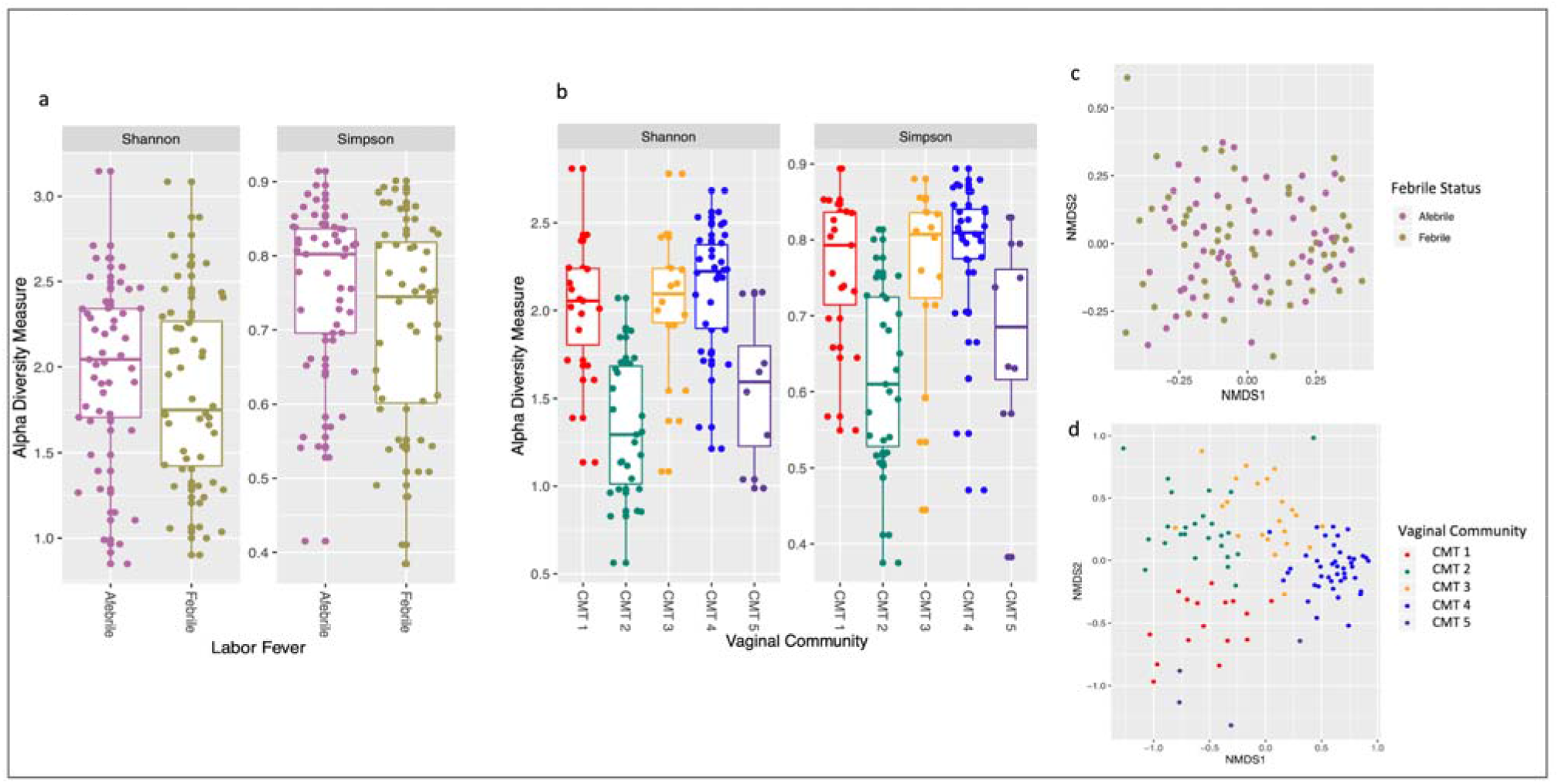
Structure of vaginal microbiome based on diversity estimates. **a**, Alpha diversity estimation (Shannon, Simpson) jitter boxplot of maternal cohort when fever status is taken into account utilizing V3-V4 regions. **b,** Alpha diversity estimation, jitter boxplot of maternal cohort when sample community assignment is taken into account; CMT denotes community (1-5). **c,** Beta diversity of maternal sample cohort shown by nonmetric multidimensional scaling (NMDS). Samples are colored based on the maternal fever status (febrile, afebrile), goodness of fit stress = 0.2. **d,** Beta diversity of maternal sample cohort by NMDS. Samples are colored by community assignment through hierarchical clustering goodness of fit stress = 0.18.

### Differential abundance analysis

We tested for associations between bacterial taxa and maternal complications, including fever status. We performed differential abundance analysis in both unadjusted and controlling for community and sample collection sites (see <u>Online Methods</u> for detail).

We observed that the abundance of G*ranucatilla, Streptococcus, Fusobacterium, Anerococcus, Sneathia, Clostridium, Gemella, Mobiluncus and Veillonella* genera were consistently higher in febrile mothers. We observed *Lactobacillus jensenii, Aerococcus spp., Prevotella copri, Acinetobacter spp., Lactobacillus crispatus and Lactobacillus reuteri* at higher abundance levels in afebrile mothers Fig. 4a and e.

**Fig. 4:**
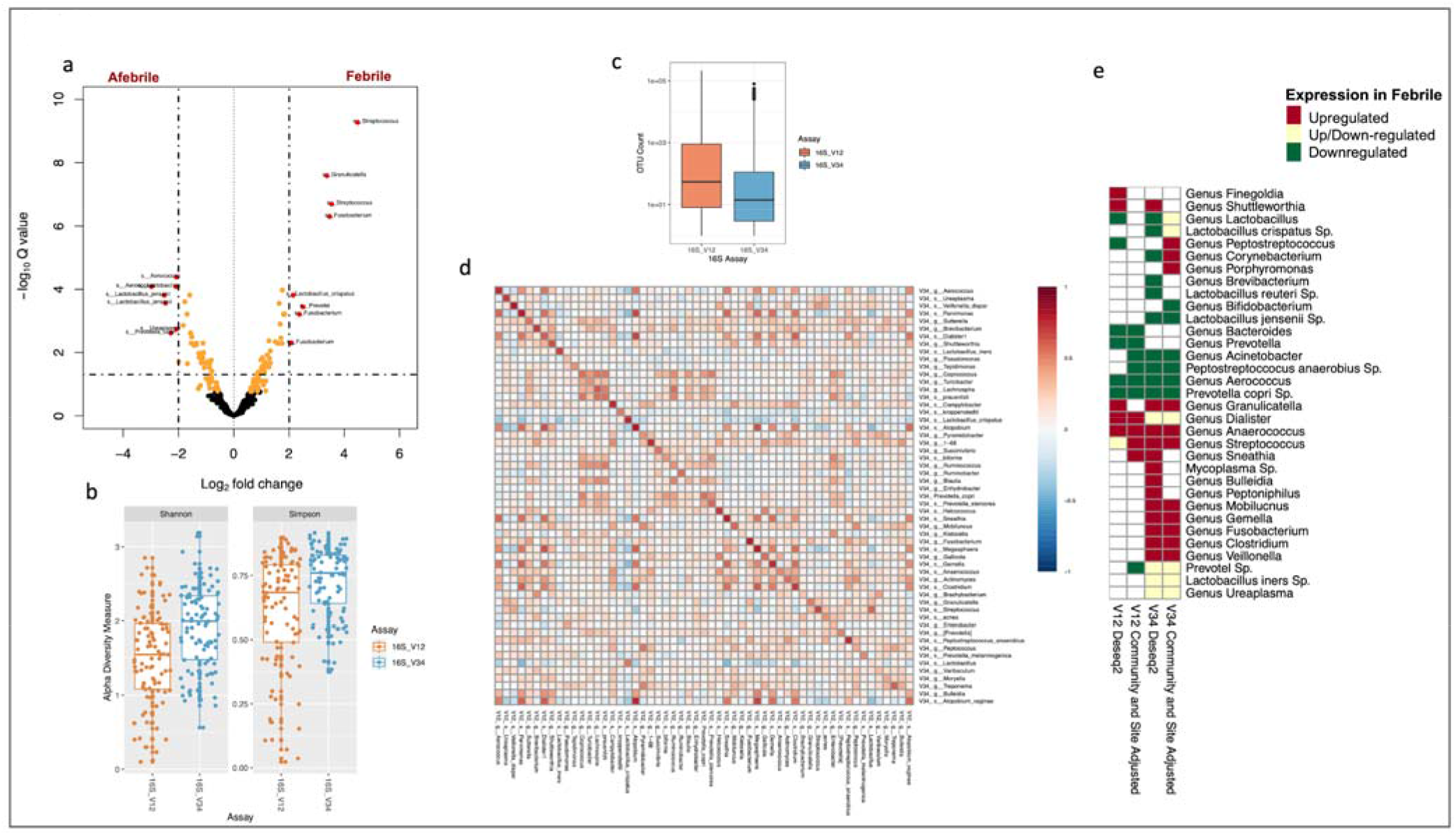
Differential vaginal bacterial presence in febrile versus afebrile laboring women utilizing V1-V2 and V3-V4 hypervariable regions. **a**, Volcano plot representing differentially expressed bacteria in febrile versus afebrile women using V3-V4 regions. Orange points signify P<0.05 and labeled red points bacteria with adjusted P<0.05 (Bonferroni correction). s_ represent species level and g_ denotes genus level classification of OTU **b,** Alpha diversity across the V1-V2 versus V3-V4 assay. **c,** Count per OTU in relevance to Assay. **d,** Spearman correlations of samples above zero counts in both V1-V2 and V3-V4 regions (OTUs agglomerated at species level). Spearman correlation measures are between (−1,1) **e,** Heatmap table of differentially expressed bacteria concordance utilizing V1-V2 and V3-V4 regions (P<0.05).

### Reproducibility with differing 16S V1-V2 and V3-V4 hypervariable regions

To assess the consistency of our results between the V1-V2 and V3-V4 hypervariable regions we compared diversity, read depth, and species abundances across matched samples. In general, alpha diversity was lower in V1-V2 relative to V3-V4 (Shannon P = 1.41e-05, Simpson P = 1.56e-4) while sample read depth was consistently higher in V1-V2 than V3-V4 (Mann Whitney P < 2.2e-16) (Fig. 4b and c).

Species abundances were highly correlated and clustered between the two hypervariable regions Fig. 4d. In particular, a cluster formed by *Turicibacter, Lachnospira, Prausnitzii, and Coprococcus* (Spearman correlation > 0.2). These bacteria are all known members of the gut microbiome family and formed a community when measuring correlation across samples. Furthermore, we identified nuanced bacterial relationships between regions. For example, *Mycoplasma spp.* displayed no correlation (Spearman correlation = 0.05) across regions; on the other hand, *Dialister* spp. had a strong correlation across both regions (Spearman correlation = 0.85) Fig. 4d. We found additional nuances between bacteria taxa depending on the hypervariable region which we have further explained in the <u>Supplementary Materials</u>.

### GoM models for characterization of vaginal microbial communities

Grades of membership models have most commonly been used in text processing and document structure identification ^31^. One such method employs a Latent Dirichlet Allocation (LDA) approach, a generative latent variable model, where documents can be thought of as bags of words generated by various themes phrased as “topics” and are represented by vectors of frequencies ^31,32^. LDA can be used to simultaneously distill topics from a set of “documents” and describe a weight to each document as belonging to a particular topic.

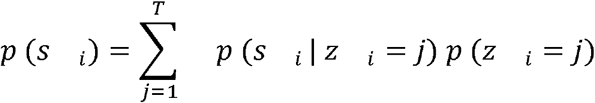

We applied GoM models to describe the underlying bacterial community structures that may have evaded detection in our hierarchical clustering methods, and to quantify the degree to which each sample belongs to the identified topics. Where *T* is the number of topics, pre-assigned through estimation of optimal numbers of clusters (i.e. topics) to be used for our GoM models via the elbow method and Nonnegative Matrix Factorization (NMF) ^33,34^. Both methods confirmed the optimal number of topics for our dataset was four (Fig. 5a, <u>Supplemental Fig. 5a</u>).

**Fig. 5:**
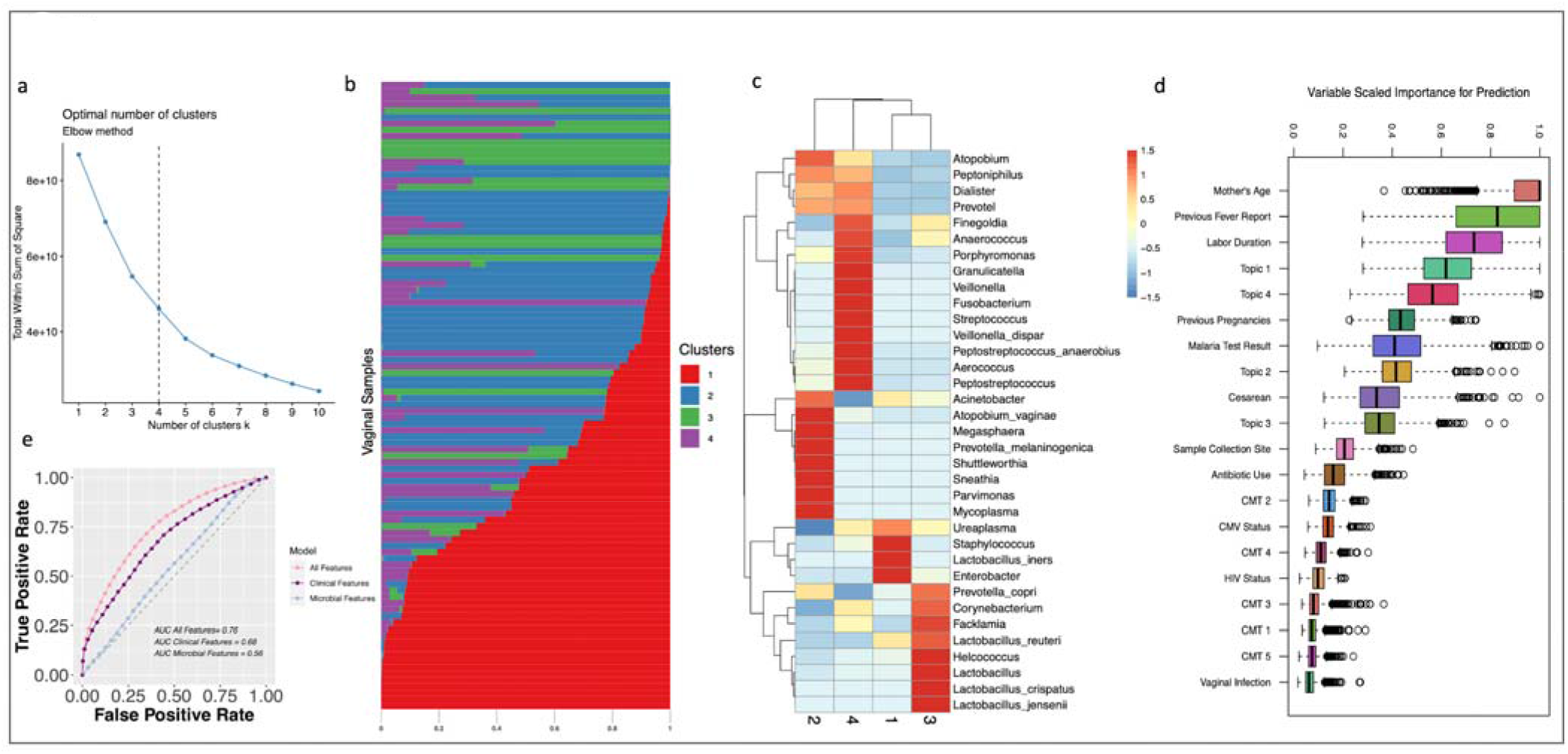
GoM models and random forest for maternal febrile status prediction. **a**, Cluster number determination for the dataset utilizing the elbow method which maximized the variance of rate of decline changes as a second derivative. **b,** The sample weight value (w) of the topic model for V3-V4 data set. Every row represents a participant’s vaginal swab sample. Every color represents the ratio in which the sample belongs to that particular cluster determined by the model. **c,** Theta (0) value for every feature (on species taxonomic rank) contributing to the formation of the clusters of GoM models. The heatmap score is a row wise z-score normalized value for every feature in each cluster. **d,** Feature importance for maternal fever status determination utilizing clinical features in addition to topic model clusters and communities identified by hierarchical clustering result of 1000 rounds of random training and test set modeling (CMT 1-5, denote communities identified in hierarchical clustering formatted as binary 1-5 (present absence feature) feature. **e,** Receiver operator curve (ROC) for 1000 rounds of random resampling of training and test set of the RF model for maternal fever status identification using all features (both clinical and microbial features), clinical, and microbial features.

Bacterial features, were agglomerated at the species level (*s*). We assume *p*(*z*) ias the distribution over topics *z* in a particular sample, this is represented as*ω*. *p*(*s* | *z*) ias the the probability distribution over bacterial species(*s*)given topic(*z*). *p*(*z_i_* = *j*) as the probability that the *j*th topic was sampled for the *i*th bacterial species and *p* (*s_i_* | *z_i_* = *j*) as the probability of the *i*th bacterial species (*s_i_*) under topic *j*, also called *θ* for simplicity. Utilizing the LDA model we determined the weight of every bacterial species in a topic(*θ*).

The key difference between the GoM models and traditional hierarchical clustering approaches include the ability to no longer rely on a dichotomous representation of a patient’s community state profile. In the hierarchical clustering a patient is explicitly given a community from the dendrogram assignment. Leveraging GoM models, we create a basis composed of a small number of microbial topics whose linear combination optimally represents each patient.

We built GoM models to evaluate the sample and bacterial distribution of our data set, and then estimated the topic weights across each sample and topic feature probability distribution, respectively (Fig. 5b, <u>Supplemental Fig. 5b</u>). We discovered a high prevalence of topics 1 and 2 across our sample set. High feature weights were observed for *Lactobacillus iners, Ureaplasma, Staphylococcus* and *Enterobacter* in topic 1. Topic 2 included *Sneathia, Shuttleworthia, Atopobium, Mycoplasma, Megasphaera* and *Parvimonas* consistently, regardless of the hypervariable region used. Topic 3 was enriched in high levels of other *Lactobacillus* species, especially *Lactobacillus crispatus, jenesenni,* and *reuteri* (Fig. 5c, <u>Supplemental Fig. 5c</u>). Topic 4 was enriched in pathogenic bacterial genera including *Granulicatella, Streptococcus, Fusobacterium, Veillonella* and *Anaerococcus* (Fig. 5c, <u>Supplemental Fig. 5c</u>, <u>Supplemental Table 5</u>).

### Clinical associations with GoM defined topics

Finally, we investigated the relationship between various clinical and environmental features of our dataset and the topics identified using our GoM model. Univariate regression defined interesting associations. Topic 1 was associated with higher odds of both prescribed and non-prescribed antibiotics use (odds ratio (OR) = 9.77, 95% confidence interval (CI) = 1.3-119.11) (<u>Supplemental Fig. 5d</u>). There was a weak correlation between increased labor duration and topic 4 (Spearman correlation = 0.20, P < 0.05). Intriguingly, topic 3, which is enriched in *Lactobacillus* species, was negatively associated with high labor duration (Spearman correlation = −0.19, P < 0.05) (<u>Supplemental Fig. 5d</u>), and topic 4 was associated with cesarean delivery (<u>Supplemental Fig. 5d</u>). Multivariate regression analysis confirmed a positive association between antibiotic use and topic 1 (P = 0.03) (<u>Supplemental Table 5</u>) when adjusting for clinical features, see <u>Online Methods</u>.

### Prediction of maternal febrile status using random forest model

To achieve a better understanding of features that may lead to intrapartum fever, we utilized a random forest (RF) classifier model, capable of handling continuous and categorical variables ^35^. We used clinical and environmental variables as features (n = 11) as well as the weights for each topic and communities identified by hierarchical clustering (CMT 1-5) (n=9). We estimated true positive and false positive rates by resampling training (n=69) and test data (n=30) from our dataset 1000 times. Maternal age was the most important feature predicting presence or absence of intrapartum fever, followed by maternal report of previous fever during pregnancy. Interestingly, topic 1, labor duration, and topics 2-4 were important correlates of maternal fever status (Fig 5d, <u>Supplemental Fig. 6b and d</u>). The performance of the model was examined using receiver operator curves (ROCs) and the area under the curve (AUC) was estimated (AUC =0.76) (Fig 5e, <u>Supplemental Fig. 6a and b</u>). Additionally, we measured the model performance for clinical and microbial features individually and found a decrease in performance when compared to the model incorporating both features (AUC=0.68 and AUC=0.56 respectively) (Fig 5e, <u>Supplemental Fig. 6b and c</u>). Further independent cohorts are necessary to validate the observed associations between these microbial topics and febrile status.

## Discussion

Heterogeneity in the vaginal microbiome could be due to environmental exposure, behavioral factors, genetic variability or medical comorbidities and may differ between continents and across populations ^36–38^. Changes in the vaginal microbial community could lead to increased risk of early-onset neonatal sepsis due to presence of pathogenic bacteria or dysbiotic communities ^37,39^. The association between the vaginal microbiome and maternal fever during labor has been incompletely studied. Here, we characterized the vaginal microbiome in laboring Ugandan women and found associations with intrapartum fever. We showed that by including novel microbial topic models we were able to predict maternal fever.

A comparison with standard microbiology versus 16S sequencing (V1-V2 and V3-V4) showed a greater sensitivity and broader bacterial capture capability utilizing 16S sequencing. Using two different hypervariable regions, V1-V2 and V3-V4, led to a more comprehensive vaginal microbiome analysis. Although commonly identified taxa were often correlated, there were unique taxa identified by each of the two regions.

We characterized the vaginal microbiome through widely-used supervised and unsupervised hierarchical clustering to identify bacteria unique to vaginal communities. As previously reported, using supervised clustering we found higher abundance of *Lactobacillus iners species* in this African population (CMT2 and CMT3) ^10^; in addition, we found higher overall diversity of bacteria across all communities compared to unsupervised approaches. The bacteria previously associated with BV (CMT IV as defined by *Ravel et al*) were both more abundant and present in more communities (CMT 2, CMT 4, CMT 5) within our cohort.

Through differential abundance analysis comparing febrile versus afebrile laboring women, we found that bacterial genera G*ranulicatella* (a nutritionally variant bacteria)*, Sneathia, Streptococcus, Fusobacterium, Clostridium* (four known BV associated bacteria)*, Anerococcus and Veillonella* (both commensal bacteria) were consistently more abundant in febrile women. Some of these genera have previously been associated with bacterial vaginosis and are known pathogens. However, to our knowledge no explicit association has been described between these bacteria and maternal intrapartum fever ^9^. The gram-negative bacterial genus *Sneathia* has been associated with high levels inflammation and the presence of vaginal proinflammatory cytokines IL-1α, IL-1β, and IL-8 proteins ^40^. High levels of group B *Streptococcus* (*agalactiae* sp.) have been described as a common cause for neonatal sepsis ^41^. Although we were unable to speciate this microbe utilizing V1-V2 primers, we were able to speciate and detect this bacteria in 29% of both febrile and afebrile patients by V3-V4 primers or bacterial recovery cultures ^41^. Previous research on group B streptococcus prevalence in pregnant women reported a 17%, 6 - 36% and 29% rate of infection in Guatemala, Denmark and the U.S respectively ^42,43^*. Lactobacillus jensenii, Aerococcus* spp.*, Acinetobacter spp., Prevotella copri, Lactobacillus crispatus, and Lactobacillus reuteri* were more abundant in afebrile women.

Leveraging the application of GoM topic models we were able to better characterize and understand the underlying structure of the antepartum vaginal microbiome. This approach enabled us to determine the best topics that fit for each individual in the cohort, rather than restricting each individual to a specific community. Through this method we identified four topics (1 to 4) prevalent in the cohort.

We identified a number of associations between the identified topics and clinical characteristics. There was an association between topic 1 and maternal antibiotic use. This association has implications for the disruption in vaginal flora that may be caused by antibiotic use, whether prescribed or self-administered. Furthermore, we found that topic 3 bacteria -- *Lactobacillus jensenii, L. crispatus, Acinetobacter spp.,* and *L. reuteri* -- abundance was associated with decreased labor duration. In contrast, topic 4 was associated with longer labor duration and cesarean delivery. Specific bacteria in topic 4 were shown more prevalent in febrile mothers, including *Streptococcus* spp.*, Granulicatella* spp.*, and Veillonella* spp. Our findings demonstrate the value of GoM topic models to categorize the structure of the vaginal microbial community in laboring women.

Finally, we fused CMT, topics, and clinical features in our data set into a random forest (RF) model. While age has been previously described as associated with febrility we found that young maternal age was the strongest factor in predicting fever. Surprisingly, topic models were shown to be among the dominant features, affirming the GoM topic approach to microbial community structure. Topics 2 and 4 were more prevalent in febrile mothers consistent with the presence of *Sneathia, Granucatilla, Anearococcus, Streptococcus* spp. in febrile women, and *Lactobacillus jensenii, L. crispatus, and L. reuteri* species in topics 1 and 3 in afebrile women.

Our findings support using a more multifaceted methodological approach leveraging multiple models to characterize such microbiomes across populations. In the case of the vaginal microbiome, this more comprehensive approach identifies structural microbiome features that better predict maternal health and risk for intrapartum women and their neonates.

## Methods

### Sample collection and ethical approval

An equal number of participants were recruited from Mbarara Regional Referral Hospital (MbararaH) in Mbarara, Uganda and Mbale Regional Referral Hospital in Mbale, Uganda (MbaleH). MbararaH is an approximately 300-bed academic hospital affiliated with Mbarara University of Science and Technology with 9,000 deliveries annually. MbaleH is a 400-bed public hospital that has nearly 10,000 deliveries a year. Both hospitals serve a mixed urban-agrarian population. Women presenting to either hospital in labor for delivery were eligible for enrollment if they were aged 18 years or older, delivered at term (≥37 weeks’ gestation), and had an intrapartum oral temperature measurement between 36.0-37.5 °C (afebrile group, *n=*25 per site); or a in intrapartum oral temperature measurement >38.1 °C on one occasion or >38.0 °C twice, at least 60 minutes apart (febrile group, *n=*25 per site).

This study protocol was approved by the institutional review boards at each participating institution, including Mbarara University of Science and Technology (MUST) Research Ethics Committee (12/11-15), Mbale Regional Referral Hospital Research Ethics Committee (082/2016), Uganda National Council of Science and Technology (HS/1963), Partners (2016P000806/PHS), and Pennsylvania State University College of Medicine (STUDY0004199).

A Materials Transfer Agreement was in place between MUST and Pennsylvania State University. An Institutional Biosafety Committee provided oversight of specimen handling at Penn State. Maternal oral temperatures were measured during labor using a digital thermometer (ADC ADTEMP Hypothermia Digital Thermometer) to confirm eligibility. After informed consent was obtained, maternal peripheral blood and vaginal samples were obtained. Maternal blood was collected using aseptic technique and tested for malaria using the rapid diagnostic test [RDT] SD Bioline Malaria Ag P.f/Pan a thick and thin blood smear. Two maternal vaginal swabs were collected consecutively during labor. First, a sterile DNA-and RNA-free swabs for 16S sequencing was inserted into the posterior vaginal fornix after gently cleansing the perineum with clean gauze. The swab rotated for 2-3 seconds, inserted into a preservative-filled (DNA/RNA Shield, Zymo Corporation) cryovial, and vortexed at high speed for 10 seconds. Swabs were then frozen in liquid nitrogen, and stored at − 80 °C prior to cryogenic transfer to the United States for sequencing. A second swab for microbiologic culture (BD BBL CultureSwab Plus Amies Gel) was then inserted into the posterior vaginal fornix immediately following collection of the first swab, rotated for 2-3 seconds and placed back into the media-containing swab container. Culture swabs collected at MbararaH were transferred to the on-site microbiology laboratory within 36 hours, while those collected at MbaleH were 4 °C for up to 48 hours before transport to a commercial microbiology laboratory in the Ugandan capital Kampala by road. Clinical data were obtained from structured interviews and chart review collected before and after delivery.

### Microbiology

Vaginal swab samples collected from MbararaH were processed in the adjacent MUST microbiology lab. A Gram stain was prepared, and swabs were plated on MacConkey agar, incubated at 37 °C, and checked for daily growth. When growth was observed, colonies were plated onto various selective and nonselective growth media and re-incubated. Following incubation, each colony type was enumerated, isolated, and identified using standard biochemical methods. Vaginal swab samples collected from MbaleH were processed at MBN Clinical Laboratories in Kampala. Interpretation of the MbararaH microbiology was performed by a single microbiologist, and interpretation of MbaleH microbiology was performed by multiple staff microbiologists.

### CMV Polymerase chain reaction (PCR)

TaqMan PCR assay targeting the cytomegalovirus (CMV) UL54 gene was utilized using previously primers and probes and PCR conditions were optimized based on recommendations by Habbal et al.^44,45^ Amplification was done on the ABI^TM^ QuantStudio 12K Real-Time PCR Instrument (USA, CA) with the following cycling conditions and times: 60°C for 30 seconds, 95°C for 5 minutes, then 45 cycles of 95°C for 15 seconds and 60°C for 1 minute. CMV positive condition was considered for samples with DNA amplification (C_t_ < 45) with technical duplicates and in cases of inconsistency a triplicate was considered. Standard curve analysis was done for all PCR runs, overall efficiency was >75%, and R^2^ was >0.95 for all runs.

### Sample extraction, library preparation and sequencing

Vaginal specimens were collected in 1 mL of DNA/RNA Shield. After collection, 1 mL of specimen was added, with swab, to 0.15 mm and 0.5 mm zirconium oxide beads and processed in a Bullet Blender (Next Advance, NY, USA) at high speed for 5 minutes. After lysis, DNA was extracted from 500 μL of the homogenized sample using ZymoBIOMICS DNA Miniprep Kit (Zymo, CA, USA) following the manufacturers protocol with proteinase K digestion and was eluted two times with 50 μL of heated elution buffer. Primer extension polymerase chain reaction (PE-PCR) of the 16S rRNA variable regions V1-2 and V3-4 was done to reduce reagent contamination using the previously described methods with region specific primers. Primer sequence are as followed: 27F: AGAGTTTGATCMTGGCTCAG, M13: CAGGGTTTTCCCAGTCACGAC, 341F_M13: CAGGGTTTTCCCAGTCACGACCCTACGGGNGGCWGCAG, and 805R: GACTACHVGGGTATCTAATCC ^46,47^. Specifically, for the V1-2 region the annealing probe was the 336R primer attached to M13 (336R_M13) and then the extended product was amplified with 27F as the downstream bacterial primer and M13. For the V3-4 region, the annealing probe was the 341F primer combined with M13 (341F_M13) and the extended product was amplified with the 805R primer and M13. Amplification was done as previously described with 500 nM primers with the MolTaq 16S Mastermix (Molzym GmbH & Co Kg, Germany) ^48^. For library preparation, after the amplified products were put into a 1x AMpure XP (Beckman Coulter) clean up, the Hyper Prep Kit (KAPA Biosystems, USA) library preparation kit was used following the manufacturer’s protocols with 7 cycles of library amplification. Library was quantified with Qubit or Agilent Bioanalyzer DNA 1000 Chip and sequenced on Illumina’s MiSeq using the 600 cycle v3 kit with 20% PhiX and 6pM pooled library. Sequencing was performed in Penn State College of Medicine Genomic core.

### Sequence alignment and taxonomic assignment

Paired-end sequences were filtered for quality using Trimmomatic (version (v) 0.38)^49^. Sequences <100 base pairs (bp) in length and average quality score <30 on a window of 20 bps were discarded. The remaining paired-end sequences were then joined utilizing PEAR v 0.9.6) ^50^. Only joined sequences with designed primers and length > 260 bp were kept. After chimeras were identified and removed using USEARCH method ^51^, sequences were clustered into operational taxonomic units (OTUs) via QIIME packages (v 1.9.1) 52. Sequences of_ over_97 percent identity represented the same genus/species, and were clustered into the same OTU, and were assigned a taxonomy by Greengenes database (v 13.8). Those OTUs without taxonomy assignment were further blasted with BlastX (v 2.7.1) to non-redundant proteins (NR) databased.

### Computational pipelines and statistical methods for downstream analysis

The biological observation matrix (BIOM) object compromising operational taxonomic unit (OTU) file, phenotype data and taxonomic assignment file was built using phyloseq and metagenomeSeq packages utilizing R programming language ^53,54^. Minimum inclusion criteria for sequenced samples was 1000 reads. Taxa with less than 2 reads in 10% of samples were excluded from the analysis. The number of Operative Taxonomic Units (OTUs) after filtering using V1-V2 primers was 274 and utilizing V3-V4 primers was 401 OTUs. For the hypervariable region comparison section taxa with fewer than 1 reads in 5% samples filtering criteria was utilized to ensure accurate estimate of presence and absence association tests for bacteria across both regions.

### Packages used for computational analysis

Beta diversity was estimated using non-metric multidimensional scaling (NMDS). Euclidean distance was measured for hierarchical clustering using the pheatmap package (v 1.0.12). Phyloseq and ggplot2 were used to estimate alpha (Shannon and Simpson) and beta (non metric multidimensional scaling) diversity and visualization of plots accordingly ^53,55^. Both adjusted and unadjusted differential abundance analyses were performed using DESeq2 ^56^. Multivariate models were adjusted for microbial community and site of collection. GoM models identified were obtained using the CountClust package^57^. Univariate and multivariate regressions were performed for topic weights regressing on various clinical features depending on the type of regression and adjusted by the number of models (i.e. based on the number of topics). Random forest (RF) models were utilized for maternal fever status prediction from the h2o package (v 3.30.0.1, https://github.com/h2oai/h2o-3).

### Statistical tests

All statistical tests and regression analyses were performed using R base functions or MASS ^58^. All P values were adjusted using Bonferroni multiple comparison test method R v 3.6.2.

## Supporting information

Main Table

## Data availability

The 16S ribosomal sequencing files and metadata could be accessed through https://microbiomedb.org/mbio/app. All the scripts and statistical methodology of this paper can be found in https://github.com/DataScienceGenomics/Vaginal?Microbiome.git.

## Acknowledgements

We are grateful to the laboring Ugandan mothers who consented to have us study their microbiomes. We thank PSU College of Medicine Genomics Sciences Core for performing all of the quality control and sequencing. Supported by an NIH Director’s Pioneer Award 1DP1HD086071 and NIH Director’s Transformative Award 1R01AI145057.

## Author Contributions

Microbiology was carried out by JB. Clinical work was performed by LMB, KB, EK, JN and MO. Data preprocessing was performed by MM and LZ. Computation and statistical analyses were performed by MM and JNP. Data management was performed by BNK and EM. Sample processing and sequencing were performed by CH and KM. Manuscript was prepared by MM, JNP and SJS. Study design was performed by MM, LMB, KB, RI, JB, JNP, and SJS. All authors contributed to editing the manuscript.

